# Orienting auditory attention in time: Lateralized alpha power reflects spatio-temporal filtering

**DOI:** 10.1101/2020.07.13.200584

**Authors:** Malte Wöstmann, Burkhard Maess, Jonas Obleser

## Abstract

The deployment of neural alpha (8-12 Hz) lateralization in service of spatial attention is well-established: Alpha power increases in the cortical hemisphere ipsilateral to the attended hemifield, and decreases in the contralateral hemisphere, respectively. Much less is known about humans’ ability to deploy such alpha lateralization in time, and to thus exploit alpha power as a spatio-temporal filter. Here we show that spatially lateralized alpha power does signify - beyond the direction of spatial attention - the distribution of attention in time and thereby qualifies as a spatio-temporal attentional filter. Participants (*N* = 20) selectively listened to spoken numbers presented on one side (left vs right), while competing numbers were presented on the other side. Key to our hypothesis, temporal foreknowledge was manipulated via a visual cue, which was either instructive and indicated the to-be-probed number position (70% valid) or neutral. Temporal foreknowledge did guide participants’ attention, as they recognized numbers from the to-be-attended side more accurately following valid cues. In the magnetoencephalogram (MEG), spatial attention to the left versus right side induced lateralization of alpha power in all temporal cueing conditions. Modulation of alpha lateralization at the 0.8-Hz presentation rate of spoken numbers was stronger following instructive compared to neutral temporal cues. Critically, we found stronger modulation of lateralized alpha power specifically at the onsets of temporally cued numbers. These results suggest that the precisely timed hemispheric lateralization of alpha power qualifies as a spatio-temporal attentional filter mechanism susceptible to top-down behavioural goals.

## Introduction

Selective attention refers to the prioritization of task-relevant sensory stimuli at the expense of distraction (Desimone & Duncan, 1995). Time and space are two fundamental dimensions across which attention is distributed. Temporal regularity in the sensory input entrains so-called “attending rhythms” (Large & Jones, 1999), which increase the attentional energy at the time points of expected stimulus occurrence and thereby improve target detection in phase with the attending rhythm (de Graaf et al., 2013; Jones et al., 2006). Furthermore, a number of recent studies found that sustained attention follows a temporally dynamic, 3-8-Hz oscillatory pattern, such that target stimuli are sampled rhythmically (e.g., Fiebelkorn et al., 2013; Landau & Fries, 2012). Besides time, space is a particularly important stimulus dimensions for object formation and selection in the visual but also in the auditory modality (Shinn-Cunningham, 2008). As an example, *spatial release from masking* describes the well-established phenomenon that target speech comprehension improves when a target talker and a concurrent masking sound become spatially separated (e.g., Arbogast et al., 2002).

A prominent signature of spatial attention in human electrophysiology is the hemispheric lateralization of the power of ~10-Hz alpha oscillations (e.g., Kelly et al., 2006; Worden et al., 2000). Across sensory modalities, alpha power in parietal and sensory regions increases in the hemisphere ipsilateral to the focus of attention and decreases in the contralateral hemisphere (audition: Ahveninen et al., 2013; vision: Bauer et al., 2012; somatosensation: Haegens et al., 2011). Relatively increased versus suppressed alpha power has been associated with the inhibition of the underlying cortical tissue (Jensen & Mazaheri, 2010) versus the release of inhibition (Strauß et al., 2014), respectively. Thus, alpha lateralization is thought of as a neural implementation of a spatial filter, which relatively increases neural processing of stimuli at relevant spatial locations and suppresses distractors. In direct support of this, we recently used an experimental paradigm to separate target selection from distractor suppression during spatial attention and found that both mechanisms independently induce lateralization of alpha power (Wöstmann et al., 2019).

Initial evidence for the behavioral relevance of lateralized alpha oscillations comes from studies that used unilateral transcranial stimulation of alpha oscillations to induce behavioral performance modulations that speak to stimulation-induced shifts in the focus of spatial attention (Deng et al., 2019; Schuhmann et al., 2019; Wöstmann et al., 2018). Besides its prominent role for spatial attention, it is less clear at present to what extent alpha lateralization also implements the dynamic distribution of attention in time.

Temporal attention can be elicited by the predictable temporal structure (e.g., its rhythmicity) of a bottom-up sensory stimulus, or by symbolic cues that provide foreknowledge about the (most likely) time point of target stimulus occurrence (for reviews on the neuroscience of temporal attention, see Herbst & Landau, 2016; Nobre & van Ede, 2018). Although there is evidence for differential neural mechanisms of temporal prediction versus attention (for review, see Schröger et al., 2015) the present study was not designed to separate independent contributions of these two. Outside of the alpha frequency range, temporal attention has been found to modulate delta (< 4 Hz) phase (e.g., Cravo et al., 2013; Wilsch et al., 2015b) and beta band activity in sensory (Todorovic et al., 2015) and in motor regions (Morillon & Baillet, 2017). Evidence from studies using multi-sensory stimuli suggest contributions of modality-specific frequency bands to temporal attention (Keil et al., 2016; Pomper et al., 2015). Important for the present study, evidence for an effect of temporal expectation on non-lateralized alpha power comes mainly from studies in the visual modality that found prestimulus alpha power modulation to reflect the anticipation of upcoming visual targets (e.g., Hanslmayr et al., 2011; Rohenkohl & Nobre, 2011; Zanto et al., 2011). Although it has been proposed that similar oscillatory processes support different types of attention (Frey et al., 2015), evidence for effects of auditory temporal attention on alpha power are more rare (Wilsch et al., 2015a).

Lateralized alpha power is modulated most prominently in-between the presentation of a spatial cue and subsequent stimulus onset and decreases thereafter (Kerlin et al., 2010; Popov et al., 2017). Nevertheless, rhythmic modulation of lateralized alpha oscillations in synchrony with the bottom-up stimulus has been observed (Kizuk & Mathewson, 2017; Tune et al., 2018; Wilson & Foxe, 2020; Wöstmann et al., 2016). These findings speak to the general sensitivity of alpha power modulation to both, bottom-up and top-down mechanisms of temporal attention.

Neuroscience work has shown that brain mechanisms of spatial and temporal attention interact. Studies that provided participants with spatial and temporal foreknowledge about visual target stimuli found overlapping parietal cortex activations in functional magnetic resonance imaging and positron emission tomography (Coull & Nobre, 1998), as well as interactive effects of both stimulus dimensions on early components in the event-related potential in the EEG (Doherty et al., 2005). In a direct attempt to test whether top-down temporal attention affects lateralized oscillatory power, van Ede and colleagues (2011) employed two different hazard rates for the occurrence of upcoming lateralized somatosensory stimuli. While lateralized power of beta oscillations (15-30 Hz) was sensitive to the type of hazard rate, the effect was weaker and not statistically significant in the alpha band. Related to this, a recent study found that temporal expectations modulate lateralized oscillatory power in a frequency- and modality-specific way (van Ede et al., 2020). It is at present unclear whether and under which circumstances the power of lateralized alpha oscillations signifies the distribution of auditory attention in time.

In the present study, we employ magnetoencephalography (MEG) to record neural responses in human participants. We ask whether and how the human brain implements an attentional filter mechanism that is both spatially and temporally specific. To this end, we augmented an established auditory spatial attention paradigm (Tune et al., 2018; Wöstmann et al., 2016; Wöstmann et al., 2018) and employed an additional cue to provide temporal foreknowledge. While pre-stimulus alpha lateralization was unaffected by temporal foreknowledge, we here show that rhythmic modulation of alpha lateralization at the speech stimulus rate increases following a cue that allows temporal foreknowledge. This modulation of alpha lateralization was temporally specific in the sense that alpha modulation was more pronounced at time points of temporally cued versus non-cued speech items.

## Materials and Methods

The design of the present study follows closely the procedure of previous auditory spatial attention paradigms used in our lab (Tune et al., 2018; Wöstmann et al., 2016). The major difference is that in addition to a spatial cue (to indicate whether to focus attention to the left versus right side), we here also employed a temporal cue to guide a listener’s attention in time to one out of five stimulus positions during a trial.

### Participants

Data of *N* = 20 participants (11 females; mean age: 26 years; range: 20-35) were included in the analysis of behavioral and MEG data. Data of one additional participant were recorded but had to be rejected due to excessive artifacts in the MEG (presumably resulting from the fact that this participant had been in an MR scanner on the previous day). None of the participants reported significant health issues or hearing deficits. Participants provided written informed consent and were financially compensated for participation. Experimental procedures were approved by the local ethics committee of the medical faculty of the University of Leipzig.

### Temporal and spatial cues

On each trial, participants received two cues providing information about the temporal position and spatial location (left vs right side) of the to-be-attended spoken number. First, they received a temporal cue, which consisted of five blue bars presented on grey background (see Fig. 1A). In one half of trials, the temporal cue was instructive, meaning that one bar was larger in height than the other bars and indicated that the respective number position was likely probed and should thus be attended. In the other half of trials, the temporal cue was neutral, meaning that all bars were of the same height and no number position was cued. For each trial with an instructive temporal cue, the number to be probed in the end of a trial was drawn from a distribution that contained the cued number positions with 70% probability and the remaining number positions with 30% probability. Instructive cue trials can be considered *valid* in case the cued number position was probed and *invalid* otherwise. The expected value of cue validity was 70%, which effectively ranged between 64% and 81% across participants in the present study.

**Figure 1.**
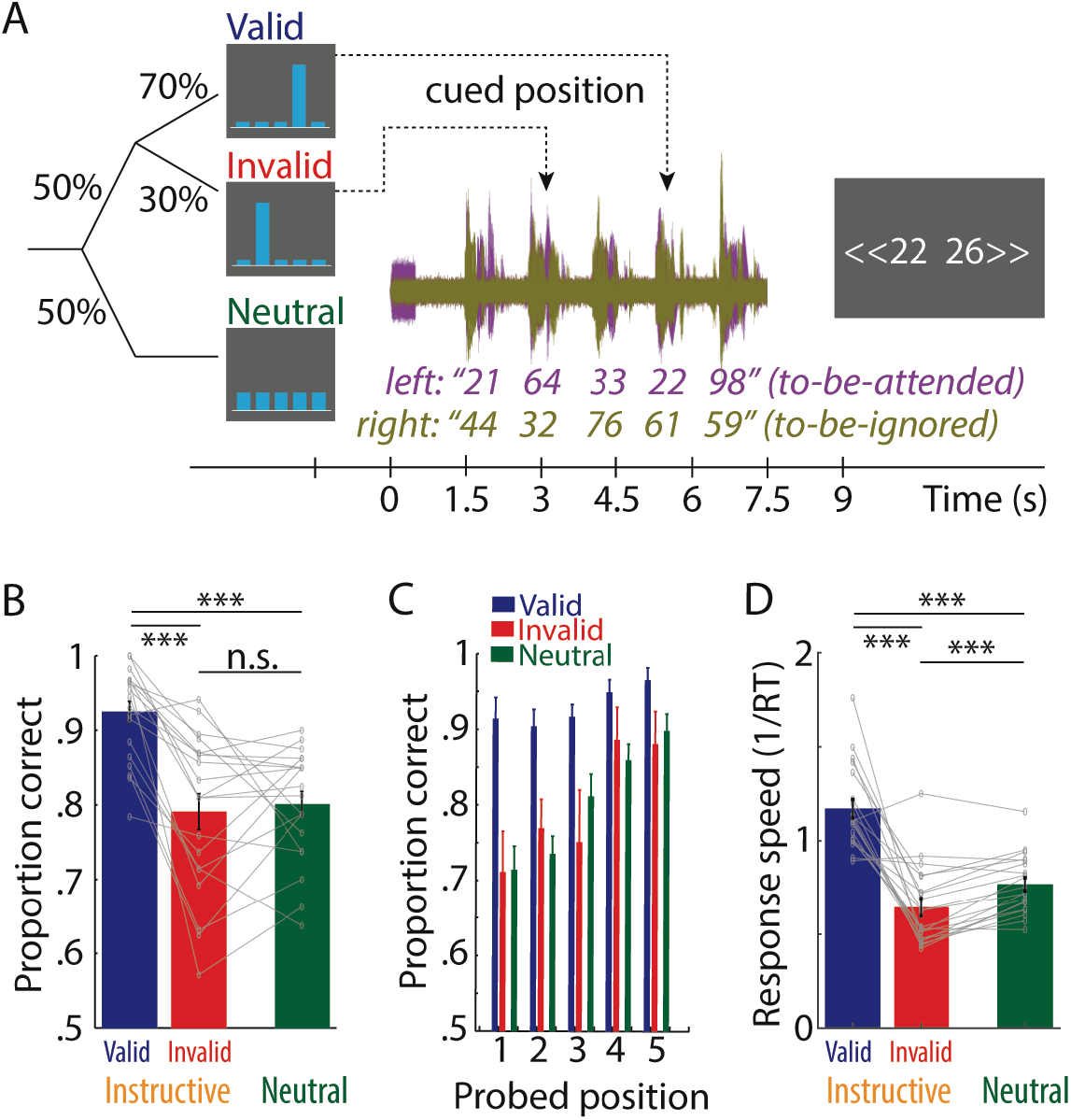
(**A**) Trial design. After a visually presented temporal cue to indicate the to-be-probed number position (valid, invalid, neutral), an auditorily presented spatial cue indicated whether spoken numbers on the left versus right side were to be attended. Participants indicated via button press which one of two numbers appeared on the to-be-attended side. (**B**) Bars and error bars respectively show average ±1 SEM proportion of correct responses for the three temporal cues. Thin lines show data of *N* = 20 individual participants. Statistical comparisons were performed on logit-transformed proportion correct scores. *** *p* < 0.001; n.s. not significant. (**C**) Same as B for individual probed number positions (1-5). (**D**) Same as B for response speed, quantified as 1/response time (RT).

Second, a spatial cue was provided to indicate whether spoken numbers on the left or right side had to be attended. The spatial cue was a monaural 1000-Hz pure tone of 0.5-s duration, with a 0.05-s linear onset ramp. The spatial cue was presented on the to-be-attended side and was valid in 100% of trials.

### Speech stimuli

Speech stimuli were the German numbers 21-99 (excluding integer multiples of ten), spoken by a female voice (adopted from Wöstmann et al., 2018). Individual numbers had an average duration (±SD) of 0.96 s ±0.05 s. Root mean square intensity was equalized across all numbers. As described in Wöstmann et al (2016), the perceptual center (P-center) of an individual number was determined (50%-point of a number’s first-syllable peak amplitude) and taken as the onset of the number for stimulus design and data analysis.

For each experimental trial, ten different numbers were randomly selected. Numbers were grouped into two streams of five numbers, each. Sound presentation was dichotic, meaning that one stream was presented to the left ear and the other to the right ear. For concurrent numbers, perceptual centers were temporally aligned, and digits were distinct in their ten and one positions (e.g., co-occurrences of “35” and “37” or “81” and “21” were avoided). The onset-to-onset interval of every two subsequent numbers was 1.25 s, resulting in a number presentation rate of 0.8 Hz. Finally, broadband background noise (at a signal-to-noise ratio, SNR, of +10 dB) was added for the entire trial duration (spatial cue onset until final number offset).

### Design and procedure

Prior to the presentation of auditory materials, the temporal cue was presented for the duration of 2.5 s and was afterwards replaced by a fixation cross. Next, after a time interval of ~1 s (jittered randomly between 0.8 and 1.2 s), the auditory spatial cue was presented either on the left or right ear to indicate that spatial attention had to be directed to the left or right side, respectively. 1.5 s after spatial cue onset, the presentation of competing numbers on the left and right side started. Approximately 1 s after the offset of the last pair of competing numbers (jittered randomly between 0.8 and 1.2 s), a response screen was presented containing two numbers. One of the probe numbers - the *target* - was a number from the to-be-attended side. The other number - the *lure* - was a number not presented at all during the respective trial. In case of a valid temporal cue trial (~70 % of instructive cue trials), the target number was the number presented on the to-be-attended side at the temporally cued position. In case of an invalid temporal cue trial (~30 % of instructive cue trials), the target number was a number presented on the to-be-attended side at a position different from the cued position. The spatial arrangement (left vs right) of target and lure on the probe screen was chosen at random on each trial. Participants used a ResponseGrip device (Nordic Neuro Lab; Norway) to indicate whether the number on the left or right was among the to-be-attended numbers with their left or right index finger, respectively. Auditory materials were presented via plastic ear tubes. Visual stimuli were shown on a back projection screen.

Each participant performed 160 trials, containing 80 trials with a neutral temporal cue and 80 trials with an instructive temporal cue. For each of these two temporal cue types (instructive and neutral), 40 trials contained a spatial cue on the left ear and 40 on the right ear. For the 80 trials with an instructive temporal cue, number positions in the middle of the trial were cued more often (positions 1 & 5: cued 8x, each; positions 2 & 4: cued 18x, each; position 3: cued 28x). Before the experiment started, participants performed 15 practice trials to become familiarized with the task. The experiment was divided in 5 blocks; participants took brief breaks in-between the blocks. The entire experimental procedure (including MEG preparation, practice trials, experiment, and a post-experiment questionnaire) took approximately 3h.

### Self-reported benefit and use of temporal cues

After the MEG experiment, participants completed a post-experiment questionnaire. They rated their benefit from instructive temporal cues (i.e. “How helpful was it for you when one blue bar was higher than the others?”; translated from German) and their use of instructive temporal cues (i.e. “How much did you use the blue bars to guide your attention to one particular number position?”) on a scale ranging from 1 (“Not at all”) to 6 (“Very much”).

### MEG recording and preprocessing

Participants were seated in a magnetically shielded room (AK3b, Vaccumschmelze). A 306-sensor Neuromag Vectorview MEG (Elekta) measured magnetic fields at 102 locations from 204 orthogonal planar gradiometers and 102 magnetometers. Here, only data recorded from gradiometer sensors were analyzed. MEG responses were recorded at a sampling rate of 1,000 Hz with a DC-300-Hz bandwidth. Each participant’s head position was monitored with five head position indicator coils. Offline, the signal space separation method (Taulu et al., 2004) was used to suppress external disturbances (i.e. noise) and transform individual participant data to a common sensor space across experiment blocks.

For subsequent MEG data analyses, we used the Fieldtrip toolbox (Oostenveld et al., 2011) for Matlab (R2018a) and customized Matlab scripts. Continuous data were highpass-filtered at 0.3 Hz and lowpass-filtered at 180 Hz, using filters described in detail in (Wöstmann et al., 2016). Data were down-sampled to 500 Hz and epochs from −2 to 10 s around spatial cue onset were extracted. Epochs were rejected when MEG responses at any gradiometer sensor exceeded 3 pT/cm. Independent component analysis was used to identify and reject components corresponding to eye blinks, saccadic eye movements, muscle activity, heartbeats, drifts, and jumps. On average, 11.75 components (SD = 4.4) were rejected.

### MEG sensor analysis

We performed two time-frequency analyses of oscillatory activity. First, for the duration of the entire trial (−2 to +10 s relative to spatial cue onset), single-trial time-frequency representations were derived using complex Fourier coefficients for a moving time window (fixed length of 0.5 s; Hann taper; moving in steps of 0.05 s) for frequencies 1 −20 Hz with a resolution of 1 Hz. Single-trial time-frequency representations were separated into four conditions, corresponding to combinations of the factors Temporal Cue (instructive vs neutral) x Spatial Cue (left vs right). Within each condition and for each one of 204 gradiometer positions, we computed oscillatory power (squaring the magnitude of complex Fourier coefficients, followed by averaging across trials) and inter-trial phase coherence (ITPC; through division of complex Fourier coefficients by their magnitudes, followed by averaging across trials). Data from 204 gradiometer sensors (102 pairs of gradiometer sensors) were then combined by averaging ITPC and by summing power estimates for every pair of sensors at the same location, resulting in estimates of power and ITPC at 102 combined gradiometers.

For further analyses, we focused on two neural measures of interest. In agreement with previous studies (Tune et al., 2018; Wöstmann et al., 2016) average low-frequency (1-5 Hz) ITPC across all 102 gradiometer sensors was taken as a neural measure of auditory encoding. As a neural index of spatial attention deployment, we calculated single-subject topographical lateralization of oscillatory 8-12 Hz alpha power (Pow) according to Equation 1:

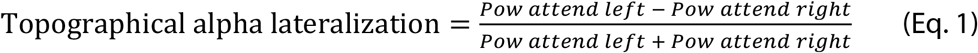

For a temporally (instead of topographically) resolved measure of alpha lateralization, we divided MEG sensors into sensors over the left versus right hemisphere (48 sensors, each). For each participant and experimental condition, 8-12 Hz alpha power was averaged across sensors on the same side as the focus of spatial attention (ipsilateral), as well as across sensors on the opposite of spatial attention (contralateral). Time-resolved lateralization of oscillatory 8-12 Hz alpha power (Pow) was calculated according to Equation 2:

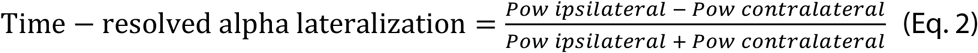

Spectral analyses of low-frequency ITPC and temporal alpha lateralization in the time window from the onset of the first number to the offset of the last number (1.5-7.5 s) was done via Fast Fourier Transform (FFT), using a Hann-window and zero-padding of the signal to achieve a frequency resolution of .02 Hz. Spectral phase and amplitude were calculated as the angle and magnitude of the complex Fourier coefficients, respectively.

Second, we analyzed alpha power lateralization for individual number positions during a trial. To this end, we split up the time-domain MEG data of each trial into sub-epochs, each ranging from −2 to +3.25 s relative to individual number onset. Note that for this analysis we did not consider the initial and the final number position on each trial, for two reasons. First, the initial and final number positions were cued very rarely (on only four trials per participant and to-be-attended side). Second, the initial number was preceded by the spatial cue and the final number was followed by behavioral response preparation, which might render MEG responses to these numbers different from the intermediate number positions 2-4.

For each participant, resulting sub-epochs around number positions 2-4 were divided into trials with spatial attention to the left versus right side. Furthermore, for trials with instructive temporal cues, sub-epochs were divided into temporally cued versus uncued. On each trial with an instructive temporal cue, there was one sub-epoch corresponding to a cued number position whereas the remaining number positions were uncued. Time-frequency representations of sub-epochs were derived using the same parameters used for the analysis of whole trial data (see above), followed by computation of oscillatory power.

By design, cued number positions were less frequent (i.e. one out of five positions in each instructive temporal cue trial) than uncued number positions (i.e. four out of five positions in each instructive temporal cue trial) and neutral number positions (i.e. all five positions in each neutral temporal cue trial). To achieve an unbiased statistical comparison of alpha lateralization for these three types of number positions, we employed a sub-sampling approach. For cued number positions, each participant’s 8-12 Hz alpha power was averaged across number positions 2-4 and trials, separately for trials with spatial attention to the left versus right side, followed by computation of time-resolved alpha lateralization (according to Eq. 2) and averaging across attend-left and attend-right conditions. For uncued and neutral number positions, we applied the same analysis on randomly drawn sub-epochs matching the quantity of cued number positions available for an individual participant, side of attention (left vs right), and number position. We repeated this step 10,000 times, to achieve 10,000 sub-samples of uncued and neutral alpha power lateralization for each participant.

For visualization purposes, alpha lateralization for all types of sub-epochs (cued, uncued, neutral) was averaged across participants, followed by overlaying of alpha lateralization for cued number positions against the 2.5^th^ and 97.5^th^ percentiles across the 10,000 sub-samples of uncued and neutral number positions. For statistical analysis, we expressed the difference of each participant’s time-resolved alpha lateralization for cued sub-epochs versus the same participant’s 10,000 sub-sampled uncued and neutral sub-epochs as z-values. This resulted in two z-values per participant and time point, one for the contrast cued vs uncued and one for the contrast cued vs neutral. For each contrast and time point, z-values were tested against zero using multiple non-parametric permutation tests (for details, see Statistical analyses). 95-% bootstrapped confidence intervals for mean z-values were computed using the bootci function in Matlab (using 10,000 bootstrap data samples).

### MEG source analysis

We used the Dynamic Imaging of Coherent Sources (DICS) beamformer approach (Gross et al., 2001) implemented in FieldTrip (version 2018-06-14). A standard singleshell head model was used to calculate leadfields for a grid of 1 cm resolution, resulting in 1,834 grid points inside the brain. To localize low-frequency ITPC, we derived a spatially adaptive filter based on the leadfield and each participant’s cross-spectral density of Fast Fourier Transforms centered at 3 Hz with ±2 Hz spectral smoothing (time period: 0-7.5 s relative to spatial cue onset). This filter was applied to single-trial Fourier Transforms (1-5 Hz, in steps of ~0.133 Hz). ITPC at each grid point was calculated and averaged across frequencies.

To localize the 0.8-Hz modulation of ITPC, the spatially adaptive filter was applied to time-resolved Fourier Transforms (centered at 3 Hz; ±0.1 Hz spectral smoothing; Hann taper; 0.5-s window moving in 0.05-s steps; time period: 0-7.5 s relative to spatial cue onset), followed by calculation of time-resolved ITPC. Spectral amplitude of 0.8-Hz ITPC modulation for each grid point was derived using an FFT (with the same parameters used for the MEG sensor analysis).

To localize alpha power lateralization prior to the onset of lateralized speech items, we first calculated a common filter for each participant, based on the leadfield and the cross-spectral density of Fast Fourier Transforms centered at 10 Hz with ±2 Hz spectral smoothing (time period: 0-1.5 s relative to spatial cue onset; calculated on all trials). Next, the common filter was used to localize alpha power separately for attend-left versus attend-right trials, followed by calculation of the alpha lateralization index for each grid point (according to Equation 1).

To localize the 0.8-Hz modulation of alpha power lateralization, a common filter was calculated for every participant with the same parameters stated above, but for the time period 1.5-7.5 s relative to spatial cue onset. The common filter was used to localize time-resolved alpha power (0.5-s window moving in 0.05-s steps; Hann taper; time period: 1.5-7.5 s) separately for four experimental conditions in the 2 (side of attention: left vs. right) x 2 (temporal cue: neutral vs. instructive) design. Since the calculation of time-resolved alpha power lateralization (according to Equation 2) requires subtraction of contralateral from ipsilateral alpha power, we first determined for every individual grid point (g_i_) the homologue grid point (g_h_) in the opposite cerebral hemisphere. Next, time-resolved alpha power lateralization for grid point g_i_ was calculated by contrasting average alpha power across g_i_ and its nine closest within-hemisphere neighbors with average alpha power across g_h_ and its nine closest within-hemisphere neighbors. Note that the degree of spatial smoothing would increase with a higher number of neighbors. Spectral amplitude of 0.8-Hz time-resolved alpha lateralization (averaged across frequencies 0.76-0.84 Hz) was determined using an FFT. Note that this procedure is only sensitive to 0.8-Hz modulation of alpha lateralization arising from the contrast of sources that are highly symmetric across the two hemispheres. Therefore, since the resulting sources of 0.8-Hz modulation of alpha lateralization are symmetric across the two hemispheres as well, we only show results for one (the left) hemisphere.

Finally, source-level ITPC and alpha lateralization were averaged across participants and mapped onto a standard brain surface. For the source-level 0.8-Hz modulation of alpha lateralization, we calculated z-values for each grid point to contrast 0.8-Hz spectral amplitude for instructive versus neutral temporal cue trials, based on dependent-samples t-tests (uncorrected).

### Statistical analyses

For statistical analyses, average proportion correct (PC) scores per participant and experimental condition (ranging from 0 to 1) were logit-transformed using an adapted logit-transform proposed by Fox & Weisberg (2019; see also Wöstmann et al., 2020).

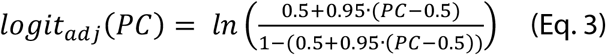

Single-trial response time (RT) data relative to the onset of the probe screen were removed in case they exceeded 3 seconds and transformed to response speed, quantified as 1/RT. Single-trial response speed data were averaged separately for individual participants and experimental conditions and submitted to statistical analyses.

As the assumption of normality was violated for some of the dependent measures in the present study, we used non-parametric permutation tests (except for circular statistics). To contrast the central tendency of two dependent variables, we calculated the relative number of absolute values of 10,000 dependent-samples *t*-statistics computed on data with permuted condition labels that exceeded the absolute empirical *t*-value for the original data (corresponding to the two-sided permutation *p*-value, denoted *p_permutation_*). To assess the relation of two variables, we calculated the relative number of absolute values of 10,000 Spearman correlation coefficients computed on data with permuted values across participants that exceeded the absolute empirical Spearman correlation coefficient for the original data (denoted *p_permutation_*).

For circular statistics, we used the Rayleigh test for uniformity of circular data and the parametric Hotelling paired sample test for equal angular means (Zar, 1999), implemented in the circular statistics toolbox for Matlab (Berens, 2009).

## Results

Twenty participants performed an adapted version of a previously used auditory spatial attention task (Fig. 1A; Wöstmann et al., 2016). On each trial, participants were presented with five spoken numbers to the left ear and - concurrently - with five competing numbers to the right ear. A pure tone in the beginning of each trial on one ear indicated the to-be-attended side. In the end of each trial, two numbers were presented on the screen. Participants had the task to select the number that had appeared on the to-be-attended side.

An ideal observer with unlimited attention and memory capacity could solve this task accurately even without temporal foreknowledge about the probed number position within a trial. Nevertheless, in order to test whether temporal foreknowledge would interact with the deployment of spatial attention, temporal foreknowledge regarding the probed number position (1-5) was triggered by a visually presented cue before each trial. This cue was either *instructive* and indicated the number position that was most likely probed (70% valid, 30% invalid) or was *neutral* (i.e. uninstructive).

### Valid temporal foreknowledge improves auditory spatial attention

For all types of temporal cues (valid, invalid, and neutral), participants performed well above chance level (proportion correct of 0.5; Fig. 1B). In the present task, any effect of temporal cues on task performance would indicate that temporal foreknowledge modifies how listeners distribute spatial attention in time. Indeed, the use of temporal foreknowledge was signified by higher proportion correct scores for valid (*Mean:* 0.93) compared with invalid (*Mean:* 0.79; statistical comparison of logit-transformed proportion correct scores, *p_permutation_* < 0.001) and neutral cue trials (*Mean:* 0.80; *p_permutation_* < 0.001). Proportion correct scores did not differ significantly for invalid versus neutral cue trials (*p_permutation_* = 0.933). This pattern of behavioral results was relatively stable across probed number positions (Fig. 1C), although overall proportion correct scores increased when later number positions were probed (test of linear coefficients for logit-transformed proportion correct as function of number position against zero; *p_permutation_* < 0.001). Measures of response speed converged with proportion correct data and revealed faster responses for trials with valid compared with invalid (*p_permutation_* < 0.001) and neutral temporal cues (*p_permutation_* < 0.001). Furthermore, response speed was lower for invalid than neutral temporal cue trials (*p_permutation_* < 0.001).

As it is typical for temporal cueing paradigms (e.g., van Ede et al., 2017), effects of cue validity in the present study were only investigated in the analysis of behavioral performance in order to verify that participants made use temporal cues. Since the validity of instructive temporal cues only became clear in the end of a trial (during probe presentation), neural responses during the trial cannot possibly be affected by cue validity. Therefore, the following analyses of MEG responses during a trial will not contrast valid with invalid temporal cues, but rather instructive versus neutral temporal cues.

### Neural signatures of auditory encoding and spatial attention

In the MEG, we focused on two neural signatures. First, low-frequency (1-5 Hz) inter-trial phase coherence (ITPC) reflected the encoding of sound events and was most prominent in temporal cortical regions (Fig. 2A). Second, the hemispheric difference in 8-12 Hz alpha power for attention to the left versus right side (i.e. alpha power lateralization) reflected participants’ deployment of spatial attention.

**Figure 2.**
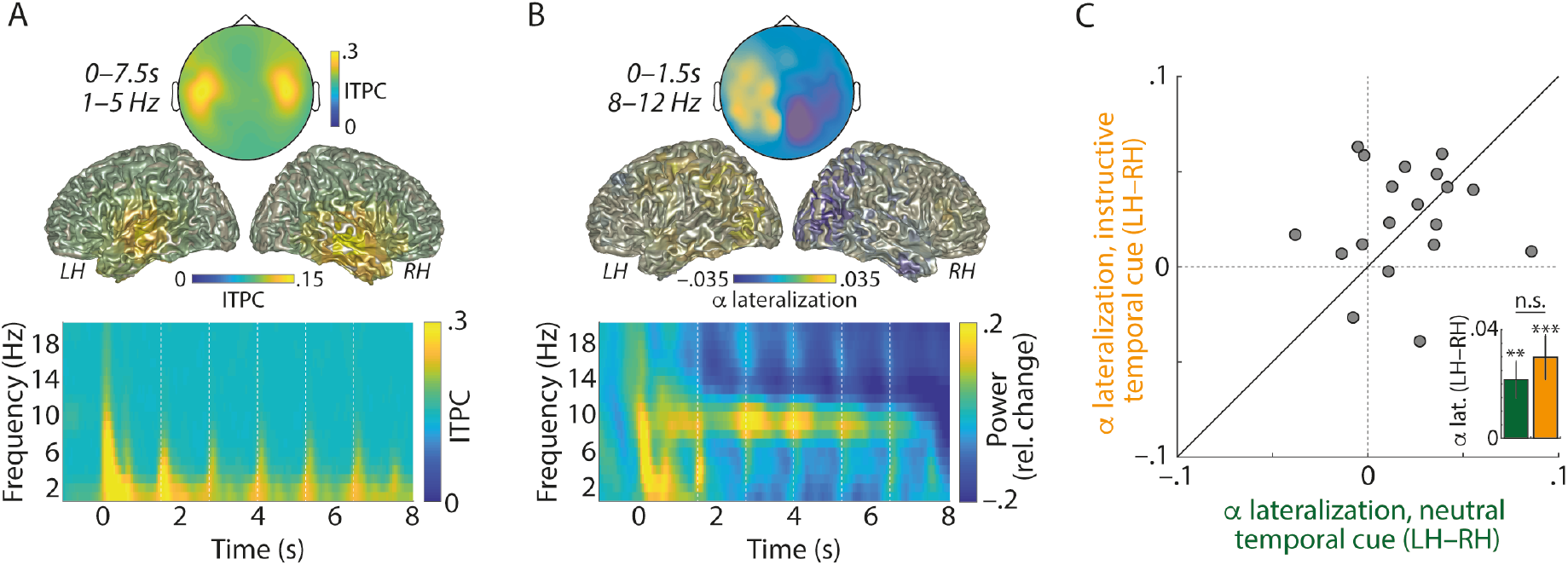
(**A**) Bottom: Grand-average time-frequency representations of inter-trial phase coherence (ITPC), averaged across all (102) combined gradiometer sensors. Dashed vertical lines indicate onsets of number positions 1-5. Top: Topographic map and brain surfaces show low-frequency (1-5 Hz) ITPC averaged across the entire trial duration (0-7.5 s). LH: left hemisphere; RH: right hemisphere. (**B**) Bottom: Grand-average time-frequency representations of oscillatory power (relative change with respect to −1 to 0 s), averaged across all (102) combined gradiometer sensors. Top: Topographic map and brain surfaces show grand-average alpha lateralization (averaged across trials with neutral and instructive temporal cues), calculated for 8-12 Hz alpha power (Pow) according to the formula: (Pow_attend-left_ - Pow_attend-right_) *I* (Pow_attend-left_ + Pow_attend-right_). The colour bar applies to both, sensor- and source-level data. (**C**) The 45-degree plot shows the hemispheric difference in alpha lateralization (LH - RH) for trials with neutral temporal cues (green, x-axis) versus instructive temporal cues (orange, y-axis). Grey dots correspond to *N* = 20 participants. Bars and error bars in the inset show average ±1 between-subject SEM of the hemispheric difference in alpha lateralization for neutral and instructive temporal cue trials. ** *p* < 0.01; *** *p* <0.001; n.s. not significant.

Before investigating the temporal modulation of ITPC and alpha lateralization during stimulus presentation, we verified whether lateralized pre-stimulus alpha power would serve as a robust measure of spatial attention, in line with previous auditory spatial attention studies (e.g., Banerjee et al., 2011; Frey et al., 2014). To this end, we calculated the topographical alpha lateralization in the time window from spatial cue onset until the onset of the first number (Fig. 2B&C; 0-1.5 s). Statistical comparison of alpha lateralization for sensors over the left versus right hemisphere revealed that alpha power was significantly lateralized (i.e. more positive values of topographical alpha lateralization on the left versus right hemisphere), for trials with neutral temporal cues (*p_permutation_* = 0.003) and for trials with instructive temporal cues (*p_permutation_* < 0.001; Supplementary Figure S1 shows that power lateralization was specific to the alpha frequency band).

Although we expected that temporal cues would affect the modulation of alpha lateralization mainly during the presentation of lateralized auditory stimuli, we also tested whether instructive versus neutral temporal cues would affect pre-stimulus alpha lateralization. This was not the case. Pre-stimulus alpha lateralization was not significantly different for instructive versus neutral temporal cues (Fig. 2C; *p_permutation_* = 0.352).

### Temporal foreknowledge enhances rhythmic modulation of lateralized alpha power

Having established neural signatures of auditory encoding and deployment of spatial attention, we next tested how instructive versus neutral temporal cues would impact the temporal dynamics of these neural signatures.

Auditory encoding, quantified as low-frequency inter-trial phase coherence (ITPC), exhibited pronounced rhythmic activity at the number presentation rate of 0.8 Hz (Fig. 3A-D). Across participants, 0.8-Hz phase angles of ITPC showed significant phase concentration for trials with instructive (Rayleigh Test; *z* = 18.23; *p* < 0.001) and neutral temporal cues (*z* = 18.21; *p* < 0.001). Neither the 0.8-Hz phase of ITPC (Parametric Hotelling Test; *F* = 0.35; *p* = 0.713) nor spectral amplitude differed between instructively-cued and neutrally-cued trials (averaged across frequencies 0.76-0.84 Hz; *p_permutation_* = 0.843).

**Figure 3.**
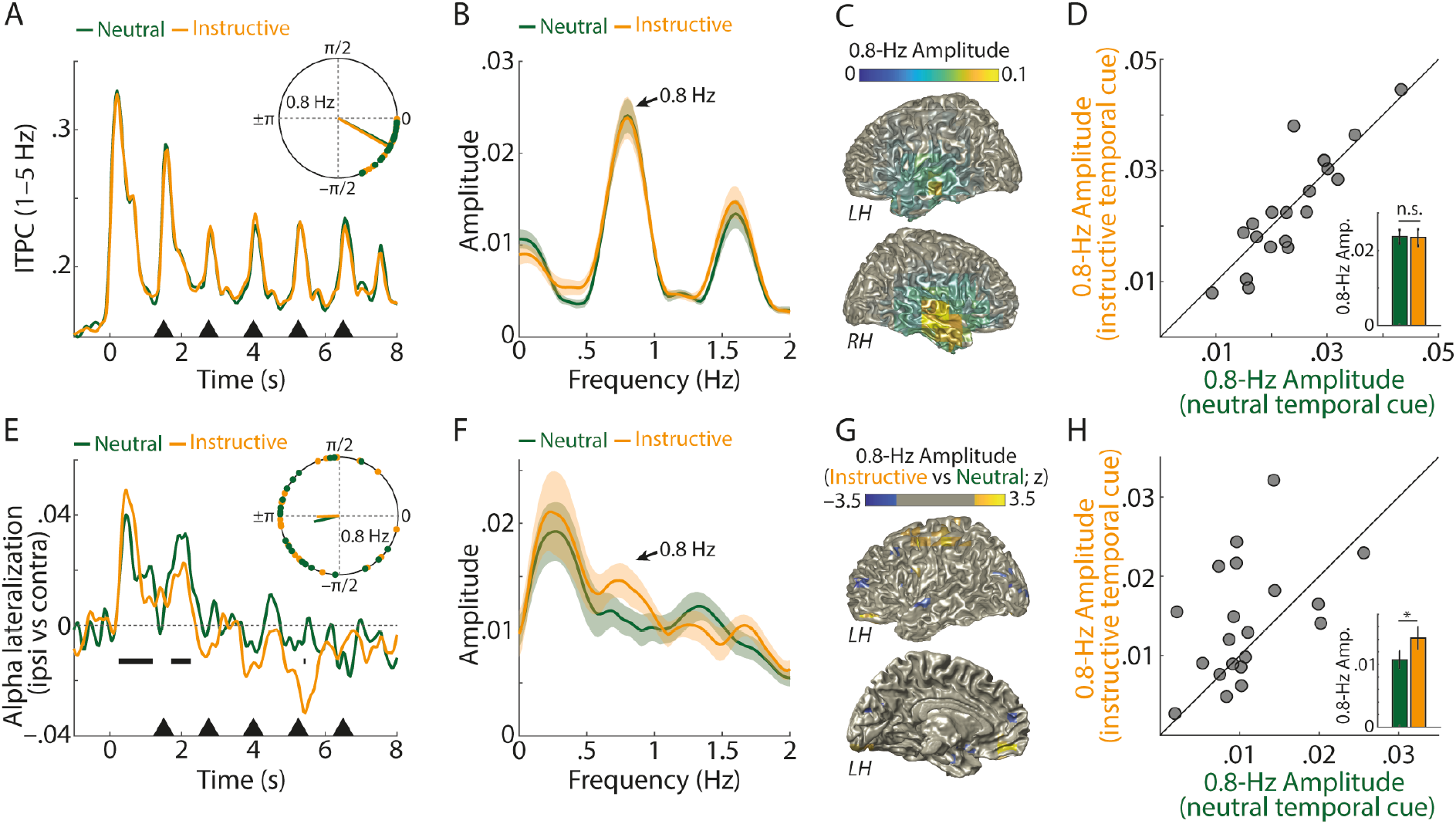
(**A**) Grand-average 1-5 Hz inter-trial phase coherence (ITPC), averaged across all (102) gradiometer sensors and *N* = 20 participants, for trials with neutral (green) and instructive temporal cues (orange). Black triangles indicate onsets of individual spoken numbers. The inset shows single-subject 0.8-Hz phase angles and resultant vectors of ITPC during number presentation (1.5-7.5 s). (**B**) Average spectral amplitude of ITPC during number presentation. Shaded areas show ±1 between-subject SEM. (**C**) Source localization of the amplitude of 0.8-Hz ITPC modulation, averaged across neutral and instructive temporal cue conditions. (**D**) The 45-degree plot shows 0.8-Hz ITPC amplitude (averaged across frequencies 0.76-0.84 Hz) for trials with neutral (green, x-axis) versus instructive temporal cues (orange, y-axis). Bars and error bars in the inset show average ±1 between-subject SEM of 0.8-Hz ITPC amplitude. n.s. not significant. (**E, F, H**) Same as (A, B, D) for alpha lateralization, calculated for 8-12 Hz alpha power (Pow) at sensors ipsilateral (ipsi) versus contralateral (contra) relative to the spatial focus of attention, using the formula: (Pow_ipsi_ - Pow_contra_) / (Pow_ipsi_ + Pow_contra_). * *p* < 0.05. Horizontal black bars in E indicate time points for which overall alpha lateralization (averaged across neutral and instructive temporal cue trials) differed significantly from zero (multiple *t*-tests; *p* < 0.05; uncorrected). (**G**) Source localization of the 0.8-Hz modulated amplitude of alpha lateralization, for instructive versus neutral temporal cue trials (z-values masked in the range: −1.96 < z < 1.96). Since the source analysis of rhythmically modulated alpha lateralization results in symmetric results in the left and right hemisphere (see Materials and Methods for details), only the left hemisphere is shown here from the outside (top) and from the inside (bottom).

The deployment of spatial attention per se, quantified as alpha power lateralization, exhibited 0.8-Hz phase concentration (Fig. 3E) for trials with neutral temporal cues (*z* = 3.84; *p* = 0.02), but the phase concentration for instructive temporal cues did not attain statistical significance (Rayleigh Test; *z* = 2.836; *p* = 0.057). Critically, the 0.8-Hz phase concentration did not differ for instructive versus neutral temporal cues (Parametric Hotelling Test; *F* = 0.12; *p* = 0.884).

Importantly, 0.8-Hz spectral amplitude of alpha lateralization was significantly increased for trials with instructive versus neutral temporal cues (Fig. 3F&H; averaged across frequencies 0.76-0.84 Hz; *p_permutation_* = 0.045). This indicates that temporal foreknowledge enhances rhythmic modulation of an established neural signature of spatial attention - the hemispheric lateralization of alpha power. This result supports our hypothesis that humans deploy alpha lateralization dynamically in time, and thus exploit alpha power as a spatio-temporal filter mechanism. The primary sources of increased 0.8-Hz modulation of alpha lateralization for instructive versus neutral temporal cue trials were found in anterior parietal cortex regions (Fig. 3G), which have previously been shown to exhibit lateralized alpha power during auditory attention (e.g. Wöstmann et al., 2019). Of note, the source localization of rhythmically modulate alpha lateralization can be considered conservative in the sense that it is only sensitive to alpha lateralization resulting from symmetric sources across the two hemispheres (see Materials and Methods for details; see also Supplementary Figure S2 for a detailed illustration of the time courses of ITPC, ipsi- and contra-lateral alpha power, as well as alpha lateralization).

It is not straight-forward to relate the rhythmic modulation of alpha lateralization to behavior in the present study, since task accuracy (to probed numbers in the end of a trial) but not neural responses to rhythmically presented speech items (during a trial) were subject to effects of temporal cue validity. However, we found that the difference in 0.8-Hz spectral amplitude of alpha lateralization for instructive minus neutral cue trials was positively related to participants’ selfreported benefit from instructive temporal cues in a post-experiment questionnaire (Fig. 4; *r_spearman_* = 0.456; *p_permutation_* = 0.044). Although the same correlation with the self-reported use of instructive temporal cues to guide attention pointed into the same direction, it was not statistically significant (*r_spearman_* = 0.425; *p_permutation_* = 0.065). These results further support the notion that 0.8-Hz modulation of alpha lateralization is part of how a listener implements temporal foreknowledge. However, it must be noted that such between-subject correlations in relatively small samples (here, *N* = 20) might be of limited robustness and should be interpreted with great care (Yarkoni, 2009).

**Figure 4.**
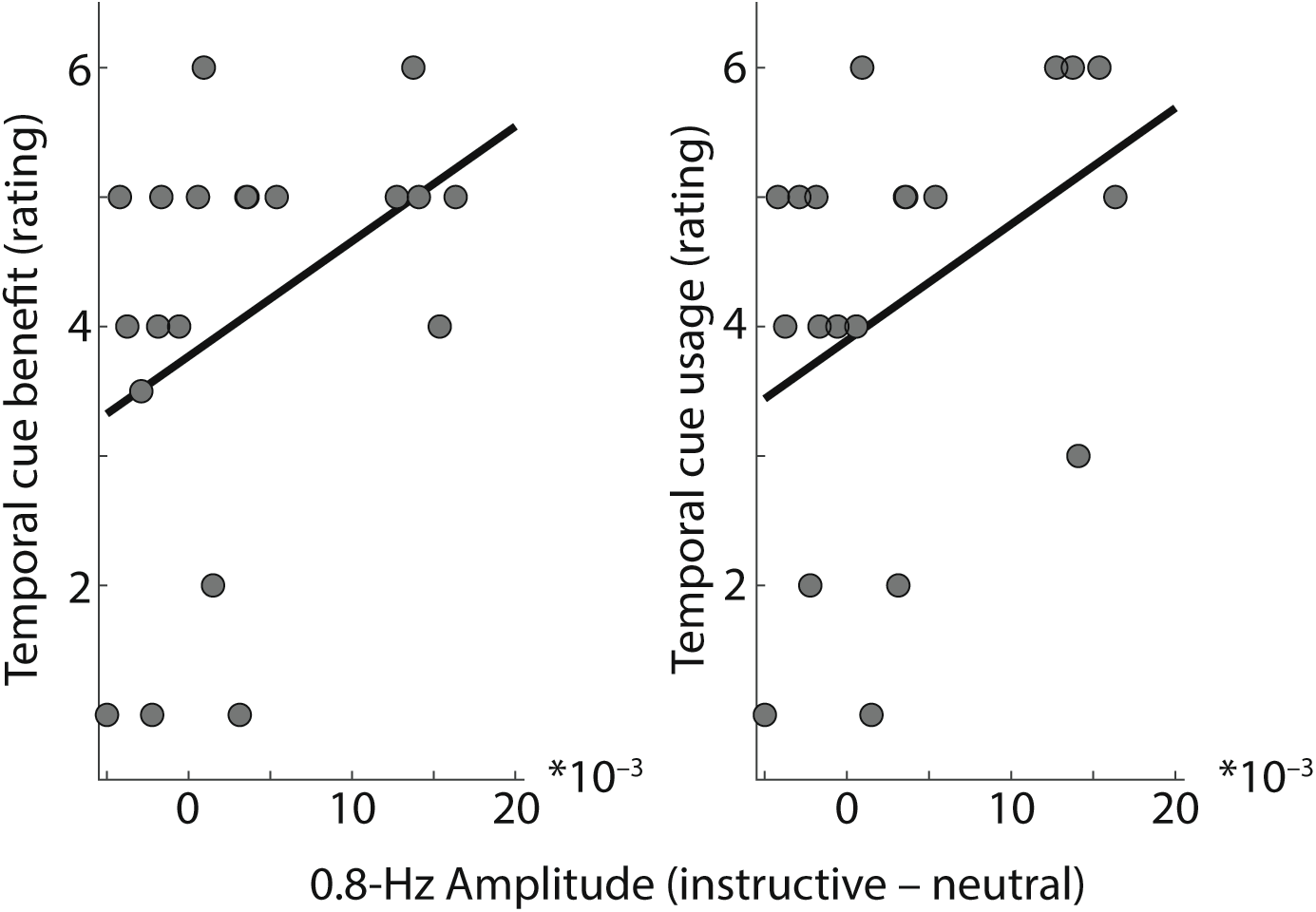
Scatterplots show relation of 0.8-Hz amplitude of alpha lateralization for instructive versus neutral temporal cue conditions (x-axis) to rated temporal cue benefit (left panel; *r_spearman_* = 0.456; *P_Permutation_* = 0.044) and temporal cue usage (right panel; *r.Spearman* = 0.425; *P_Permutation_* = 0.065).

### Lateralized alpha modulation by foreknowledge is temporally specific

It is of note that the analysis of rhythmic modulation of lateralized alpha power across the whole time window of number presentation (Fig. 3) is of limited temporal specificity, since instructive temporal cues only cued one particular (out of five) number positions within a trial. For this reason, we next analysed the modulation of lateralized alpha power for individual number positions, which were *cued* or *uncued* in instructive temporal cue trials and *neutral* in neutral temporal cue trials (see Supplementary Figure S3 for an analysis of ITPC for individual number positions).

Figure 5 shows the contrast of alpha lateralization for cued versus uncued number positions and for cued versus neutral number positions (Supplementary Figure S4 shows that the lateralization of oscillatory power around number onsets was specific to the alpha frequency band). It might at first glance seem surprising that we did not observe more positive, but instead more negative alpha lateralization for cued versus uncued numbers (Fig. 5C) and for cued versus neutral numbers (Fig. 5D) in a ~0.5-s long time window including number onset. However, as we know from the whole-trial analysis (Fig. 3), alpha lateralization temporally lags the presentation and sensory encoding of numbers by ~0.46 s. Thus, there is a more negative state of alpha lateralization at number onset, followed by a state of alpha lateralization close to zero ~0.46 s thereafter. For cued compared with uncued and neutral numbers, the difference between these two states is more pronounced, which effectively results in a stronger modulation of lateralized alpha power.

**Figure 5.**
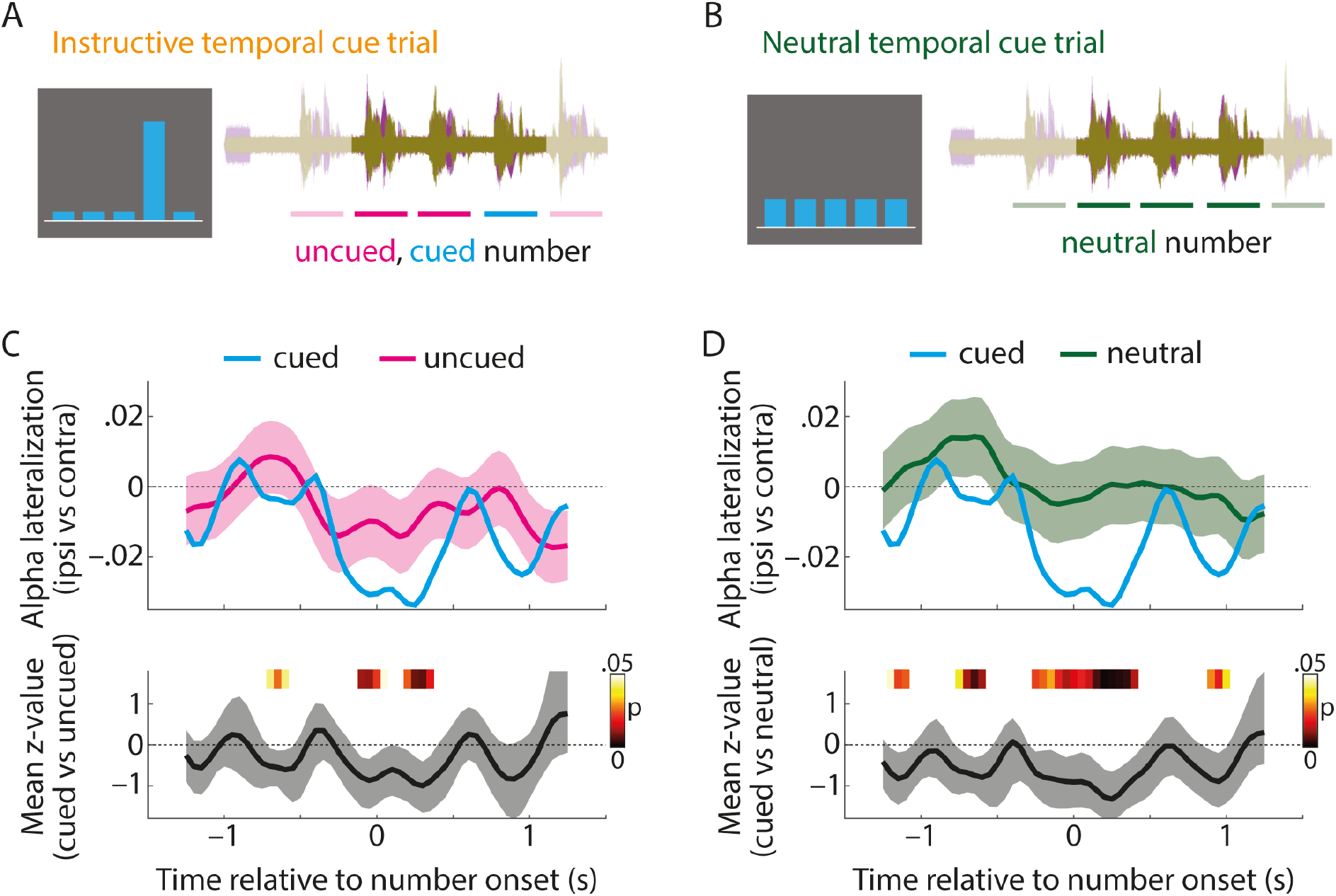
(**A**) In each trial with an instructive temporal cue, there were four temporally uncued and one cued number positions. (**B**) In each trial with a neutral temporal cue, all five number positions can be considered neutral. Note that the first and last number positions are shaded in A&B, since these number positions were not considered for the present analysis (see Materials and Methods for details). (**C**) Top: Temporal alpha lateralization index for cued number positions (blue) and uncued number positions (pink; calculated by sub-sampling the same quantity of uncued MEG epochs as there were cued epochs available for each participant, number position, and side of attention; see Materials and Methods for details). Shaded areas show the 2.5^th^ and 97.5^th^ percentiles of alpha lateralization across 10,000 sub-samples of uncued number positions. Since no sub-sampling was required for cued number positions (see Materials and Methods for details), no error bars are shown. Bottom: Solid line and shaded area show the average across single-subject Z-values to quantify alpha lateralization for cued versus uncued number positions and 95-% bootstrap confidence interval, respectively. P-values (indicated in color) were derived by testing Z-values of *N* =20 participants against zero using multiple non-parametric permutation tests. (**D**) Same as C but for the comparison of cued versus neutral number positions.

## Discussion

The present study was designed to test whether temporal foreknowledge modulates the spatial attention-driven hemispheric lateralization of alpha power in a dichotic listening task. The most important results can be summarized as follows. First, higher accuracy and response speed for valid versus invalid and neutral temporal cues indicate that participants use temporal foreknowledge to guide spatial attention. Second, temporal foreknowledge increased the rhythmic modulation of lateralized alpha power, which demonstrates sensitivity of the lateralized alpha response to spatial but also temporal attention. Third, modulation of lateralized alpha power at the onset of cued versus uncued temporal positions speaks to the temporal specificity of alpha power modulation in spatio-temporal filtering.

### Lateralized alpha responses in auditory spatial attention

Although evidence for lateralization of alpha power in spatial attention tasks is most abundant in the visual (e.g. Kelly et al., 2006; Worden et al., 2000) and somatosensory modalities (e.g. Bauer et al., 2012; Haegens et al., 2011), spatial attention to auditory stimuli induces lateralized alpha power as well (e.g. Ahveninen et al., 2013; Banerjee et al., 2011). Since left and right auditory cortices tend to receive input primarily from the contralateral ear (Rosenzweig, 1951; Tervaniemi & Hugdahl, 2003), a straight-forward functional interpretation is that relatively high versus low alpha power in one hemisphere suppresses versus enhances processing of the input to the contralateral ear, respectively.

Although alpha oscillators have been identified in auditory cortex regions (Billig et al., 2019; Keitel & Gross, 2016; Lehtelä et al., 1997) and although lateralized alpha responses during auditory spatial attention have been localized in auditory cortex regions (Müller & Weisz, 2012; Weisz et al., 2014; Wöstmann et al., 2016), alpha lateralization is typically most pronounced in parietal cortex regions, which host a supramodal attention network (Banerjee et al., 2011). In agreement with this literature, lateralized alpha power following spatial cues in the present study was strongest in parietal regions (Fig. 2B), which might speak to a shift of supramodal spatial attention rather than auditory attention alone. In general, imaging modalities with a high spatial resolution (such as Electrocorticography, ECoG; de Pesters et al., 2016) or sophisticated study designs that allow separation of auditory from visual and somatosensory alpha responses (e.g. Spitzer et al., 2014) are necessary to identify alpha responses originating from auditory cortex regions (for a review on alpha oscillations in audition, see Weisz et al., 2011).

Temporal foreknowledge in the present study had no significant impact on rhythmic auditory encoding (quantified by ITPC) but on the rhythmic deployment of spatial attention (quantified by lateralized alpha power). This is in general agreement with previous studies using a similar paradigm (Tune et al., 2018; Wöstmann et al., 2016), which found that low-frequency ITPC is mainly driven by the bottom-up stimulus presentation and does not serve as an index of attention deployment.

### Temporal evolution of the lateralized alpha response

Although the main goal of the present study was to investigate the fluctuation of lateralized alpha power in synchrony with the speech rate (0.8 Hz), the more global temporal evolution of lateralized alpha power (Fig. 3E) deserves attention as well. In general, positive values of alpha lateralization reflect the well-established pattern of higher ipsilateral than contralateral alpha power during spatial attention (e.g. Kelly et al., 2006; Worden et al., 2000), which has been proposed to reflect the attentional selection of targets and suppression of distractors (Wöstmann et al., 2019). In theory, negative values of alpha lateralization (such as they show up later during a trial in Fig. 3E and in Fig. 5) are unexpected, since these would reflect the opposite, that is, selection of distractors and suppression of targets.

However, there are three possible reasons why the sign of the alpha lateralization index later during a trial should be interpreted with caution. First, it must be noted that there are striking differences in early (prestimulus) and later (peristimulus) alpha power lateralization: While the former is almost exclusively positive in spatial attention tasks, the latter has been observed to exhibit values close to zero or even marginally negative (Kerlin et al., 2010; Tune et al., 2018; Wöstmann et al., 2016). One possible interpretation is that early alpha lateralization reflects a trial-initial shift of spatial attention to a relevant position, whereas later alpha lateralization signifies smaller relative adjustments of the established focus of spatial attention.

Second, since the alpha lateralization index is a (normalized) difference metric, its signal-to-noise ratio (SNR) is lower than the SNR of the original time courses of ipsi- and contralateral alpha power (the same issue is well-known in the domain of difference waves of event-related potentials; see e.g. Luck, 2005). In the present study, the confidence interval of the alpha lateralization index further increased later during a trial and included zero for the vast majority of time points after 2 s (Supplementary Figure S5).

Third, a recent study suggests that late (peristimulus) but not early (prestimulus) alpha lateralization decreases in case non-spatial features aid separation of auditory streams: Alpha lateralization during the stimulus period was virtually nullified when a large pitch separation aided allocation of attention to one auditory stream in space (Bonacci et al., preprint). Small or even negative values of alpha lateralization later during the trial in the present study might thus have several reasons. Trials in the present study were relatively long and participants presumably established a spatial focus of attention mainly in the beginning of the trial after spatial cue presentation. Furthermore, the availability of non-spatial, i.e. temporal, information in the present study might have reduced the overall reliance on spatial cues. For these reasons, we consider it more important to focus on the pattern of rhythmic fluctuations rather than the overall level of lateralized alpha power.

In fact, our results suggest that the timing of relatively more positive versus more negative states of lateralized alpha power is crucial when it comes to the interpretation of the functional role of lateralized alpha power for spatio-temporal filtering. In agreement with what we have observed in previous studies, which used the same auditory spatial attention task without temporal cues (Tune et al., 2018; Wöstmann et al., 2016), the average 0.8-Hz modulation of alpha lateralization lagged behind the average 0.8-Hz modulation of ITPC by ~0.46 s (corresponding to a 133° phase lag). This indicates that the auditory encoding of two competing numbers was followed, after ~0.46 s, by a relatively more positive alpha lateralization. A necessary by-product of a more positive lateralized alpha response after ~0.46 s is a more negative lateralized alpha response at the time of number onset (Fig. 5). Presumably, the temporally lagging alpha lateralization is a neural signature of a reactive filter mechanism, which serves to select the previously presented number on the to-be-attended side and to filter out the number on the to-be-ignored side (for an overview of different mechanisms of attentional filtering, see Geng, 2014). In agreement with the lagging time course of alpha lateralization in the present study (see also Fig. S2), relatively delayed lateralized alpha responses have also been observed in other multi-talker scenarios (e.g. Getzmann et al., 2020). Although direct evidence for a reactive filtering account of lateralized alpha power is missing, two recent studies in the visual modality at least speak against a purely proactive role of lateralized alpha power for sensory gain control (Antonov et al., 2020; Gundlach et al., 2020).

In addition to effects of spatio-temporal attention deployment, we cannot exclude that participants learned the statistical regularities of the distribution of task-relevant stimuli, which occurred more often at intermediate than at earlier or later number positions. Related to this, it has been proposed that prior experience with distracting information might shape the neural processing thereof (for a recent review, see van Moorselaar & Slagter, 2020), even in case the statistical regularities are unknown to the observer (Wang & Theeuwes, 2018). Future studies would have to implement systematic variation of the distribution of task-relevant stimuli to test whether and to what extent learned statistical regularities contribute to the observed temporally specific modulation of lateralized alpha power.

### Limitations

Although we found converging behavioural and neural evidence for an effect of temporal foreknowledge on auditory spatial attention, the observed effects on the neural level are of limited size. We consider four main reasons for this. First, our instructive temporal cues had a validity of only 70%. Despite the fact that a certain number of invalid cues are required in order to empirically test whether participants made use of temporal cues, higher cue validity should result in stronger and more temporally-specific allocation of temporal attention. Second, although valid temporal cues clearly improved participants’ performance, temporal cue usage was not strictly required to perform correctly in the present task. Third, an analysis of temporal cueing effects on individual number positions 1–5, which might in principle provide further insights on the temporal dynamics of spatial attention deployment, was not possible in the present study due to a very limited number of trials per cued number position. Fourth, the conventional pattern of positive alpha lateralization in spatial attention tasks was absent (and even marginally negative) later during the relatively long trials in the present study, which arguably compromises the interpretation of the temporal modulation of alpha lateralization as a direct signature of the attention filter.

Despite the straight-forward interpretation of alpha lateralization lagging auditory encoding (Fig. 3), the present results cannot completely reject the possibility that the lateralized alpha response instead precedes auditory encoding. In order to contrast proactive versus reactive accounts of filtering, future studies could decouple stimulus presentation and participants’ expectation thereof to disentangle whether alpha power lateralizes prior to expected stimulus onsets or following actual stimulus onsets. Furthermore, it is at present unclear in how far the results of the present study translate to sensory modalities other than audition. A recent study that employed low-frequency ITPC and lateralized alpha power in the MEG (Wilsch et al., 2020) found that effects of temporal expectation are stronger in audition, whereas effects of spatial attention are stronger in vision. Thus, the presence and size of interactive effects of temporal and spatial attention might depend on the sensory modality under investigation.

## Conclusion

The present data move us beyond previous research that had shown non-lateralized alpha power to be modulated by temporal expectation more generally (Rohenkohl & Nobre, 2011; Wilsch et al., 2015a), and lateralized beta power to be modulated by temporal hazard during spatial attention (van Ede et al., 2011). The present study demonstrates that the power of lateralized alpha oscillations - a well-established index of spatial attention - signifies the distribution of attention in time. In the light of previous research where behavioural benefits were accompanied by stronger modulation of lateralized alpha power (Wöstmann et al., 2016), an obvious interpretation is that temporal foreknowledge increases the temporally specific modulation of lateralized alpha power to enhance the spatio-temporal filtering of task-relevant auditory information.

## Data availability statement

All data are available from the corresponding author upon request.

## Acknowledgements

This research was funded by an ERC consolidator Grant (ERC-CoG-2014-646696 AUDADAPT to J.O.).

---

**Figure S1.**
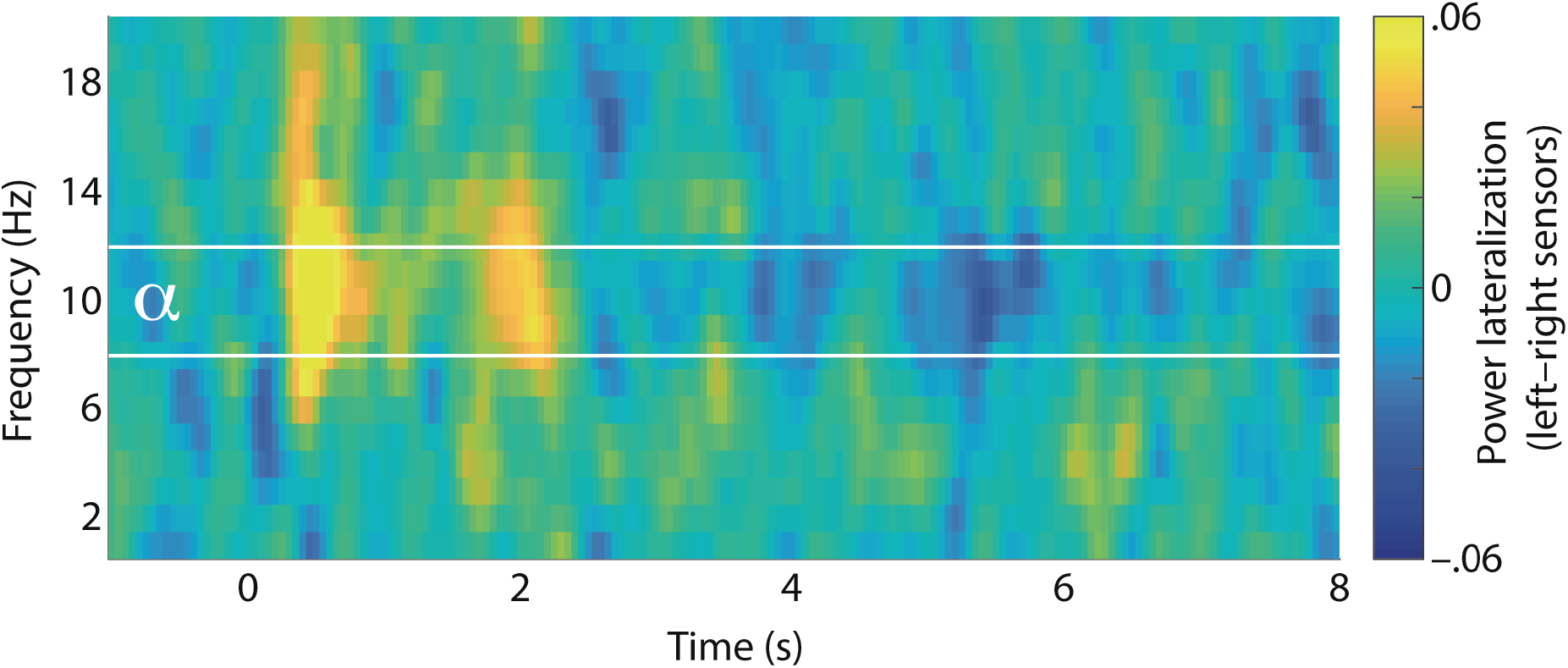
Time-frequency representation of the lateralization index calculated on oscillatory power (Pow) according to the formula: (Pow_attend-left_ − Pow_attend-right_) / (Pow_attend-left_ + Pow_attend-right_) for all temporal cue conditions (neutral and instructive). The index was calculated at all sensor positions, followed by averaging across left- and right-hemispheric sensors separately, and calculation of the left-minus right-hemisphere difference. Accordingly, positive values indicate higher ipsilateral than contralateral oscillatory power. Hemispheric lateralization of power was strongest in the first third of a trial (~0-2.5 s) in the alpha frequency range (8-12 Hz; highlighted by white horizontal lines), but decreased thereafter and eventually became negative (in agreement with Figure 3E in the main manuscript).

**Figure S2.**
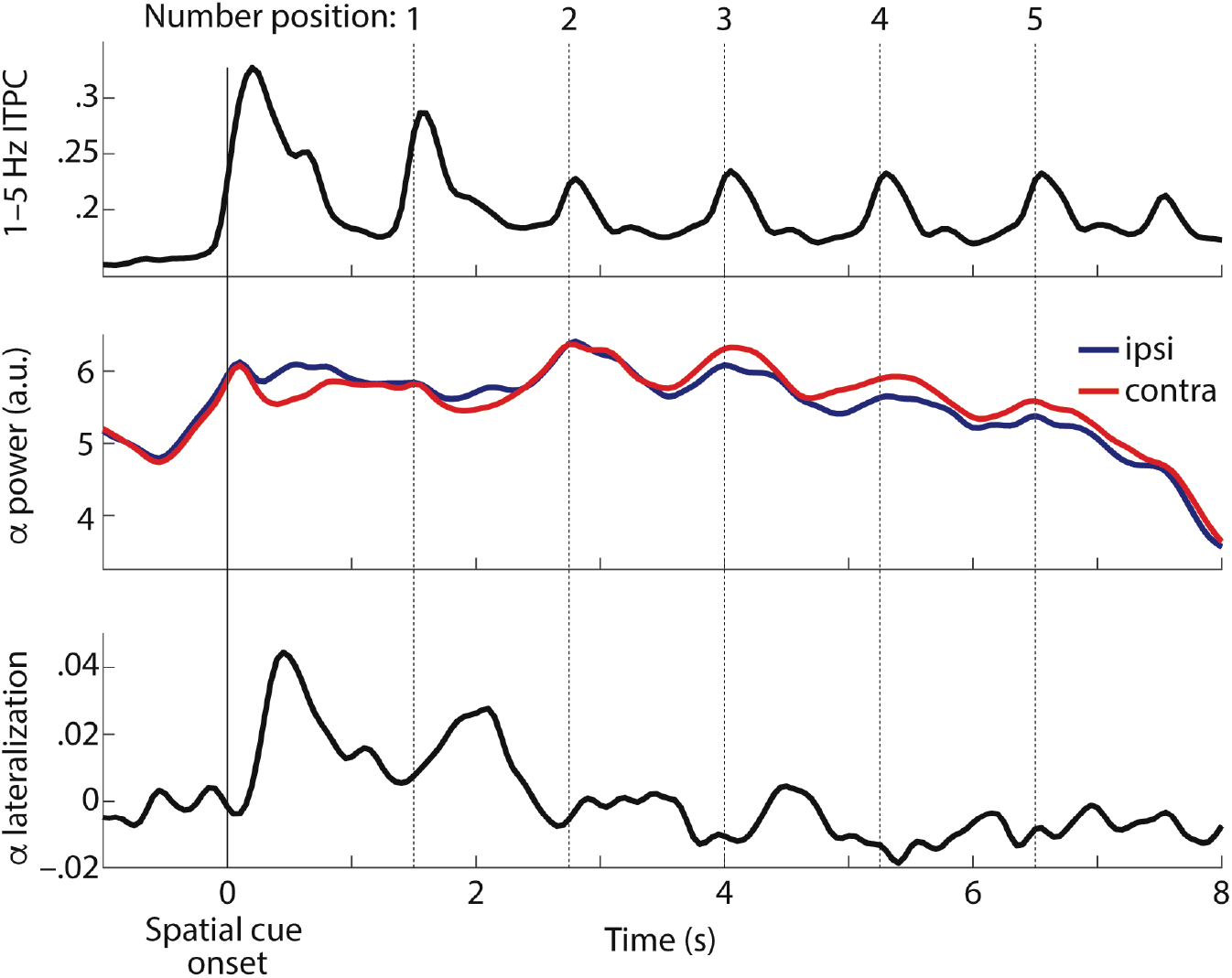
Top: Time course of low-frequency ITPC (averaged across all *N* = 20 participants, temporal and spatial cue conditions, all 102 combined gradiometer sensors, and frequencies 1-5 Hz). Middle: Time course of absolute alpha power ipsilateral (blue) and contralateral (red) to the focus of spatial attention (averaged across all *N* = 20 participants, temporal and spatial cue conditions, all 102 combined gradiometer sensors, and frequencies 8-12 Hz). Bottom: Time course of the alpha (8-12 Hz) lateralization index, calculated on alpha power (Pow) at sensors ipsilateral (ipsi) versus contralateral (contra) relative to the spatial focus of attention, using the formula: (Pow_ipsi_ − Pow_contra_) / (Pow_ipsi_ + Pow_contra_). Alpha lateralization was averaged across all *N* = 20 participants, and both temporal cue conditions. Vertical solid and dashed lines indicate the onset of the spatial cue and numbers, respectively.

**Figure S3.**
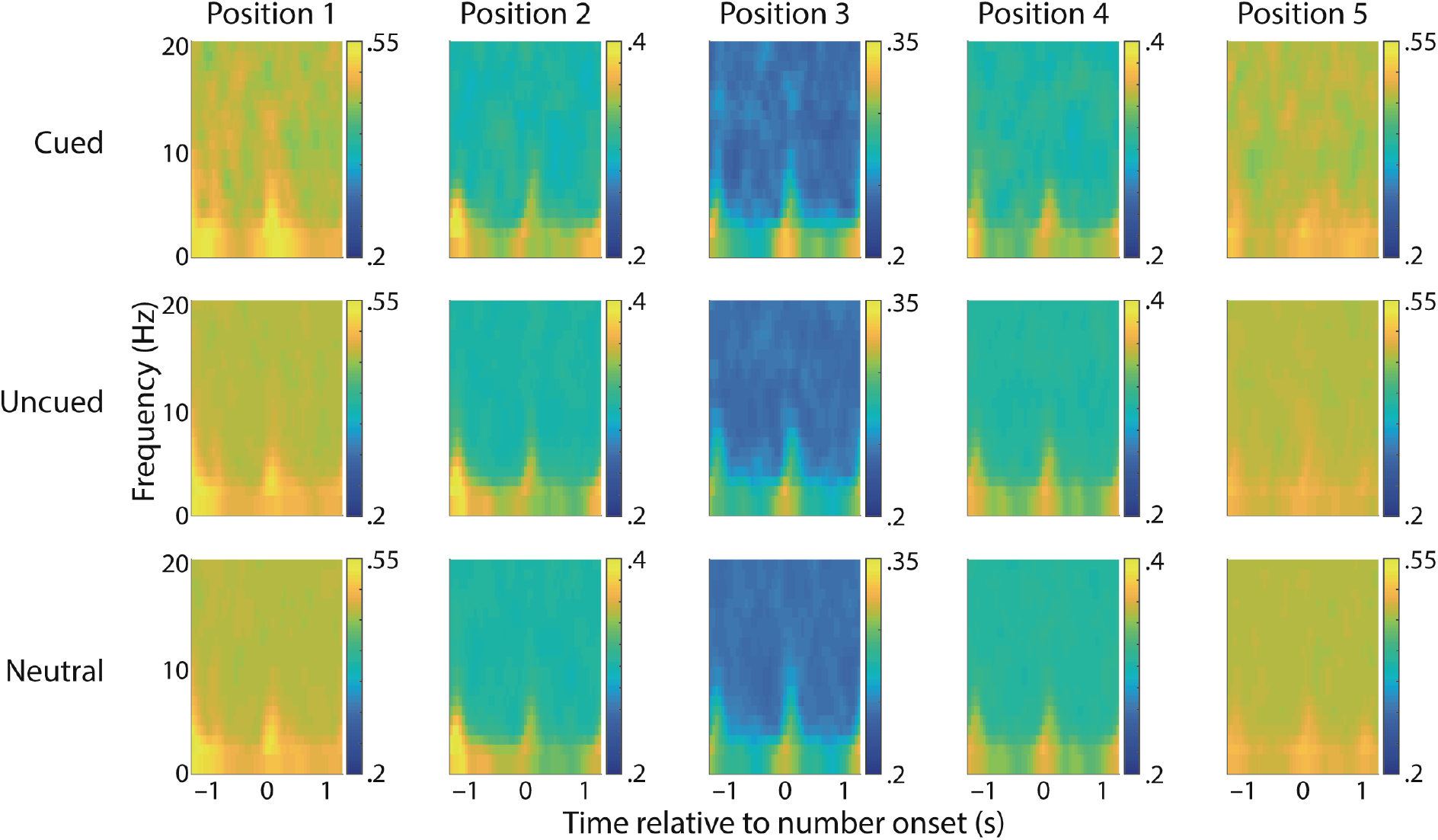
ITPC for individual number positions (columns) and temporal cueing conditions (rows). ITPC was averaged across all (102) combined gradiometer sensors and *N* = 20 participants. By design, there were less cued than uncued number positions and neutral number positions. For this reason, a subsampling procedure was used that selected for each participant, attended side, and number positions as many uncued and neutral trials as there were cued trials, at random without replacement. This procedure was repeated ten times and ITPC was calculated and averaged across the ten subsamples. Since ITPC critically depends on trial number, ITPC is generally higher for positions with fewer trials (i.e. positions that were cued less often). However, the sub-sampling procedure allows to compare ITPC across temporal cueing conditions (i.e. within each column). Note different scaling of colour bars.

**Figure S4.**
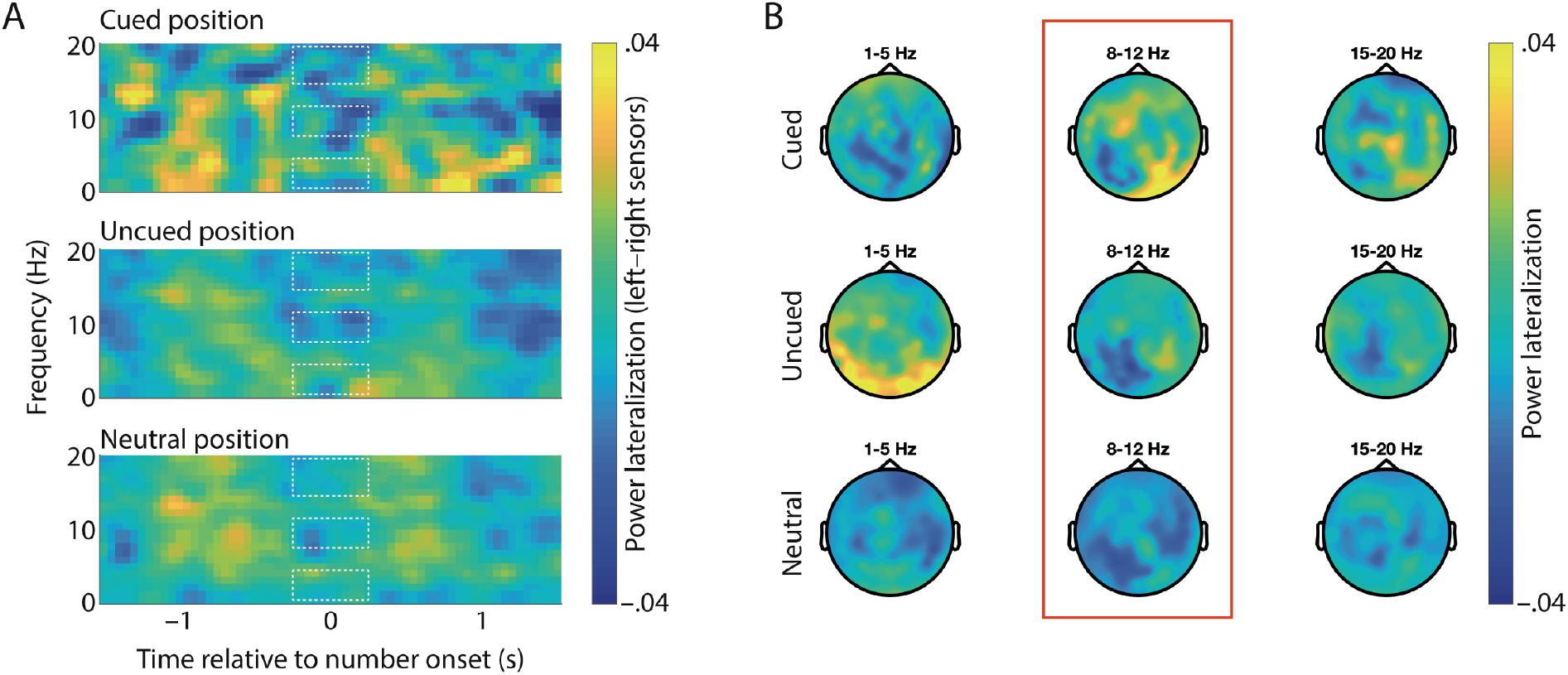
(**A**) Same as Figure S1, but for shorter time intervals around cued (top), uncued (middle), and neutral number positions (bottom) within a trial. Positive values indicate higher ipsilateral than contralateral oscillatory power, and vice versa for negative values. Dashed boxes indicate the time interval −0.25 to +0.25 around number onset in the theta (1-5 Hz), alpha (8-12 Hz), and beta (15-20 Hz) frequency bands. (**B**) Topographic maps show power lateralization for different cueing conditions (rows) and frequency bands (columns), calculated according to the formula: (Pow_attend-left_ - Pow_attend-right_) *I* (Pow_attend-left_ + Pow_attend-right_). In agreement with what is shown in Figure 5 in the main manuscript, alpha lateralization was more negative (i.e. smaller values of the lateralized alpha response on the left versus right hemisphere) in the alpha band at the onset of cued compared with uncued and neutral positions (highlighted by red box). This effect was clearly less obvious in the other frequency bands and can thus be considered alpha-specific.

**Figure S5.**
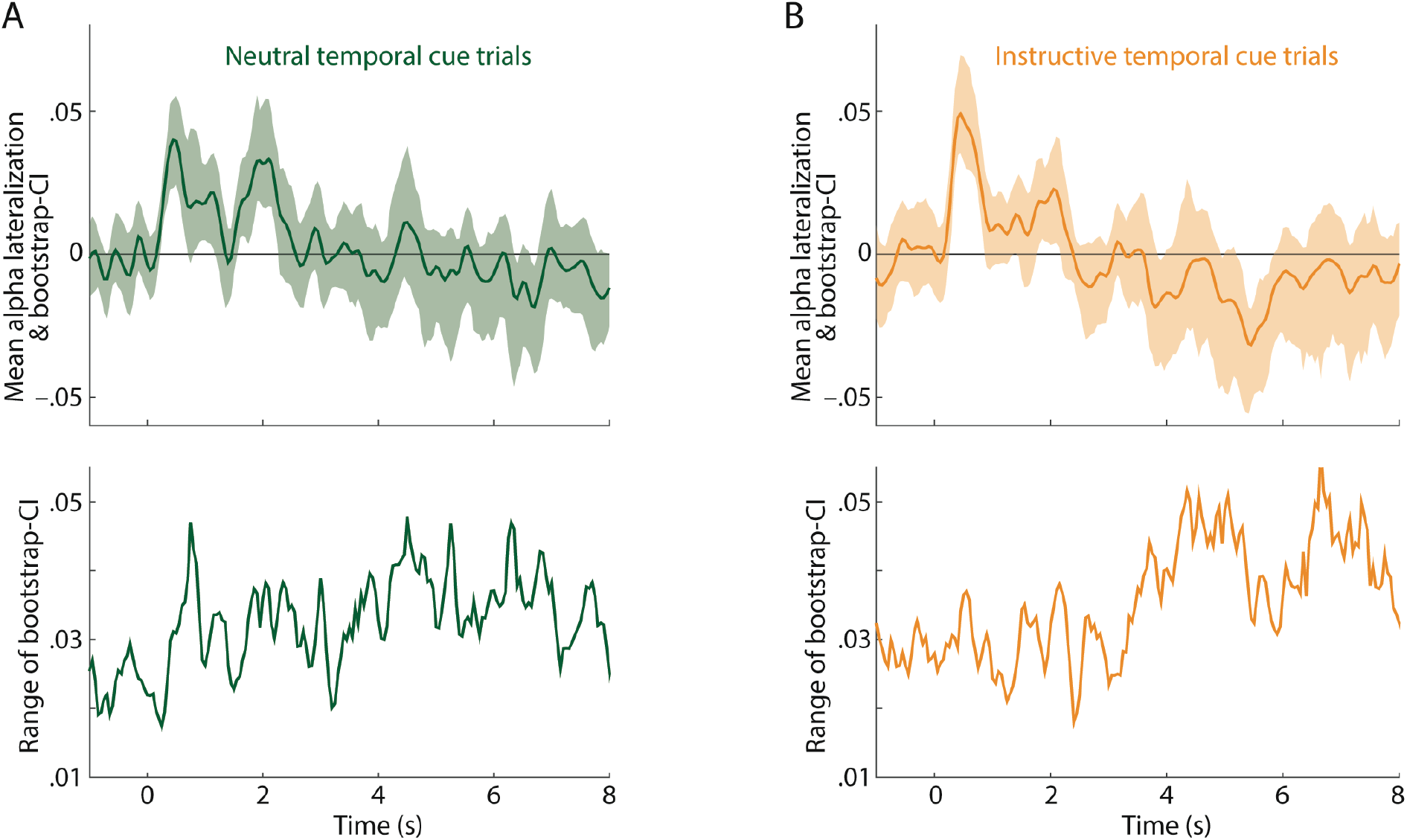
(**A**) Top: Solid line and shaded area show the mean alpha lateralization index (similar to Figure 3E in the main manuscript) for neutral temporal cue trials and 95-% bootstrap confidence interval (CI; computed using 1,000 bootstrap samples in the *bootci* function for Matlab), respectively. Bottom: Range of the bootstrap-CI as a function of time. (**B**) Same as (A) but for instructive temporal cue trials. Note that the range of the bootstrap-CI tends to increase later during a trial, which supports the claim that the sign of the late lateralization index should be interpreted with caution.

